# THE *HOK/SOK* TOXIN/ANTITOXIN LOCUS ENHANCES BACTERIAL SUSCEPTIBILITY TO DOXYCYCLINE

**DOI:** 10.1101/2020.02.13.948752

**Authors:** Chinwe U. Chukwudi, Liam Good

## Abstract

The antibacterial efficacy of the tetracycline antibiotics has been greatly reduced by the development of resistance, hence a decline in their clinical use as antibiotics. The *hok*/*sok* locus is a type I toxin/antitoxin plasmid stability element, often associated with multi-drug resistance plasmids, especially ESBL-encoding plasmids. It enhances host cell survivability and pathogenicity in stressful growth conditions, and particularly increases bacterial tolerance to β-lactam antibiotics. The *hok*/*sok* locus forms dsRNA by RNA:RNA interactions of the toxin and antitoxin, and doxycycline has been reported to bind and inhibit dsRNA cleavage/processing. This study investigated the antibacterial efficacy of doxycycline in hok/sok host bacteria cells, the effect on *hok*/*sok*-induced growth changes and the potential mechanism of the observed changes. Different strains of *E. coli* with growth characteristics affected by the *hok*/*sok* locus were transformed with *hok*/*sok* plasmids, and assessed for doxycycline susceptibility and growth changes. The results show that the *hok*/*sok* locus increases bacterial susceptibility to doxycycline, especially in strains with more pronounced *hok*/*sok* growth effects. The increased doxycycline susceptibility occurs despite β-lactam resistance imparted by *hok*/*sok*. Doxycycline was found to induce bacterial death in a manner phenotypically characteristic of Hok toxin expression, suggesting that it inhibits the toxin/antitoxin dsRNA degradation, leading to Hok toxin expression and cell death. In this way, doxycycline could be used to counteract the multi-drug resistance plasmid maintenance/propagation and pathogenicity mechanisms associated with the *hok*/*sok* locus. This has great potentials in the global war to contain the rise in antimicrobial resistance.

## 1 Introduction

Many cellular RNAs play a regulatory role, and the secondary structures that are important to their biological activities are formed through folding and formation of double-stranded regions (dsRNA). Antisense RNAs are examples of regulatory RNAs, which appear to act mainly by down-regulating gene expression post-transcriptionally following the formation of dsRNA with their target messenger (sense) RNAs. By forming a duplex with the mRNA of the regulated gene via base-pairing interactions, the dsRNA regions may block ribosome recognition of the mRNA or lead to recognition and cleavage by double-stranded ribonucleases [1, 2]. In bacteria, many antisense RNAs are encoded on plasmids, and are often involved in regulation of plasmid replication and copy number control [3]. Many plasmid and chromosomally-encoded antisense RNAs in bacteria are known to act as type I antitoxins by down-regulating the expression of toxic proteins [4-6]. In addition, some of these antisense RNA antitoxins are known to regulate the expression of toxins associated with stress response [4].

A well-established example of *cis*-encoded regulatory antisense RNA in bacteria is the Sok (<u>s</u>uppression <u>o</u>f <u>k</u>illing) antitoxin commonly found in *E. coli*. The Sok antisense RNA regulates the expression of the *hok* (<u>ho</u>st <u>k</u>illing) mRNA by binding to the *hok* mRNA and initiating RNAse III decay of the duplex, thereby inhibiting the translation of the *hok* transcript [7, 8]. Sok RNA is more labile and easily degraded (half-life of about 30s) than the stable *hok* mRNA (half-life of hours), which persists for much longer within new daughter cells following cell division [9-11]. In daughter cells containing the plasmid, the continued rapid transcription of Sok antisense RNA from its strong promoter ensures inhibition of *hok* expression. In cells that lose the plasmid during division, the acquired Sok RNA are more rapidly degraded than the *hok* mRNA, thus releasing the stable *hok* mRNA for translation, leading to subsequent cell death by the toxin produced (Fig 1). The toxin causes damage to the cell membrane that appears as the characteristic morphology referred to as ‘ghost cell’ [11-14]. This ensures that all surviving daughter cells inherit the plasmid, as those that did not receive the plasmid at replication are quickly killed by the Hok toxin [15]. Additionally, the *hok*/*sok* locus has been found to enhance bacterial stress response and improve growth in growth-limiting conditions such as high temperature, low cell density and antibiotic treatment [16].The R1 plasmid in which the *hok*/*sok* locus was originally discovered carries several genes that encode resistance traits, and in this way is able to impart multi-drug antibiotic resistance (ampicillin, chloramphenicol, kanamycin, streptomycin and sulphonamides) to its host bacteria [17-19]. Subsequently, the *hok*/*sok* locus has been found in many plasmids that encode extended spectrum beta-lactamases, especially CTX-M ESBLs [20], as well as in the chromosomes of some enterobacteria [21]. In *E. coli*, the chromosomally-encoded *hok*/*sok* loci are more abundant in pathogenic strains [22]. In laboratory strains (K-12), some chromosomal loci appear to be inactivated by insertion elements located close to the toxin reading frames, which are absent in wild type cells [4]. Unlike the plasmid-encoded locus, the chromosomally-encoded *hok*/*sok* loci do not mediate plasmid stabilization, and their *hok* toxin mRNAs are poorly translated *in vitro* [21]. Hence, it has been suggested that translational activation may require induction by unknown signals in living cells.

**Figure 1:**
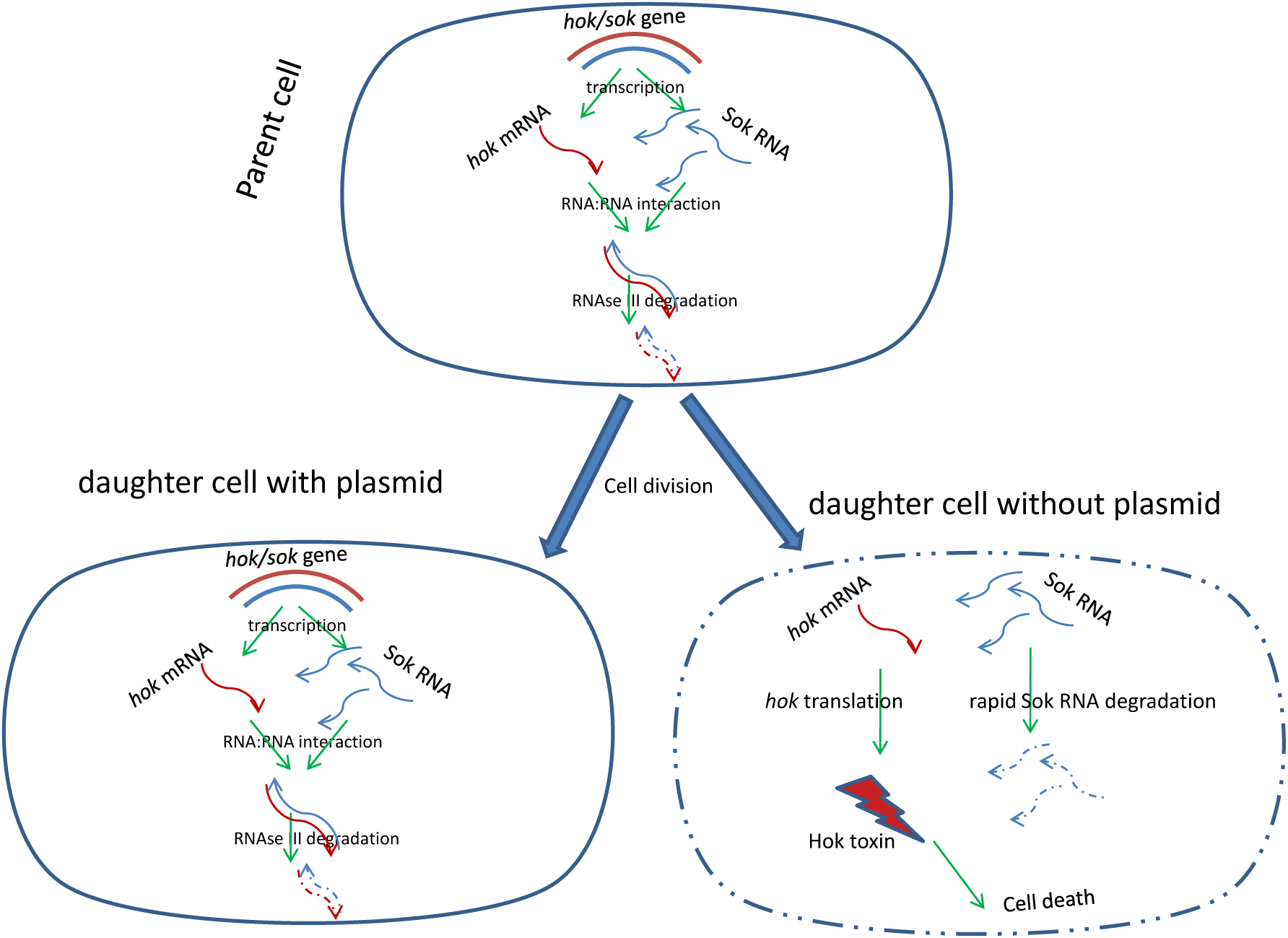
Schematic presentation of the *hok*/*sok* locus as RNA:RNA interaction model involved in plasmid stabilization via post-segregational killing. In cells containing the *hok*/*sok* plasmid, Sok (<u>s</u>uppression <u>o</u>f <u>k</u>illing) antitoxin RNA forms dsRNA with the *hok* (<u>ho</u>st <u>k</u>illing) toxin mRNA which are degraded by RNAse III to prevent the expression of the toxin. In the case of plasmid loss (e.g. following cell division), Sok RNA is rapidly degraded, releasing the *hok* mRNA for translation. The Hok toxin released produces ‘ghost cell’ morphology and cell death.

Doxycycline has been shown to bind dsRNAs and inhibit their cleavage by RNAse III [23]. It has also been shown to inhibit pre-rRNA processing and the formation of mature rRNA in *E. coli*; a pathway primarily dependent on RNAse III cleavage [24]. Hence, we hypothesize that doxycycline could inhibit RNAse III degradation of the *hok*/*sok* toxin/antitoxin dsRNA complex, leading to the release of the *hok* mRNA for translation as the labile Sok RNA disintegrates, and consequent cell death. Given that the *hok*/*sok* locus is often associated with antibiotic resistance genes and propagation mechanism, this study investigated the effect of doxycycline on *hok*/*sok* host bacteria survivability and associated antibiotic resistance mechanism.

## 2 Materials and methods

### 2.1 Strains and plasmids

*E. coli* strains and plasmids used in this work are listed in Table 1. Cells were grown on LB agar/broth, with the addition of 100µg/ml of ampicillin (Amp) or 30µg/ml of chloramphenicol (Chlr) where indicated. Antibiotic susceptibility disks were purchased from Thermo Fisher Scientific (Oxoid). Bacteria stocks were stored in 15% glycerol at -20°C for short periods and −80°C for long periods.

**Table 1:**
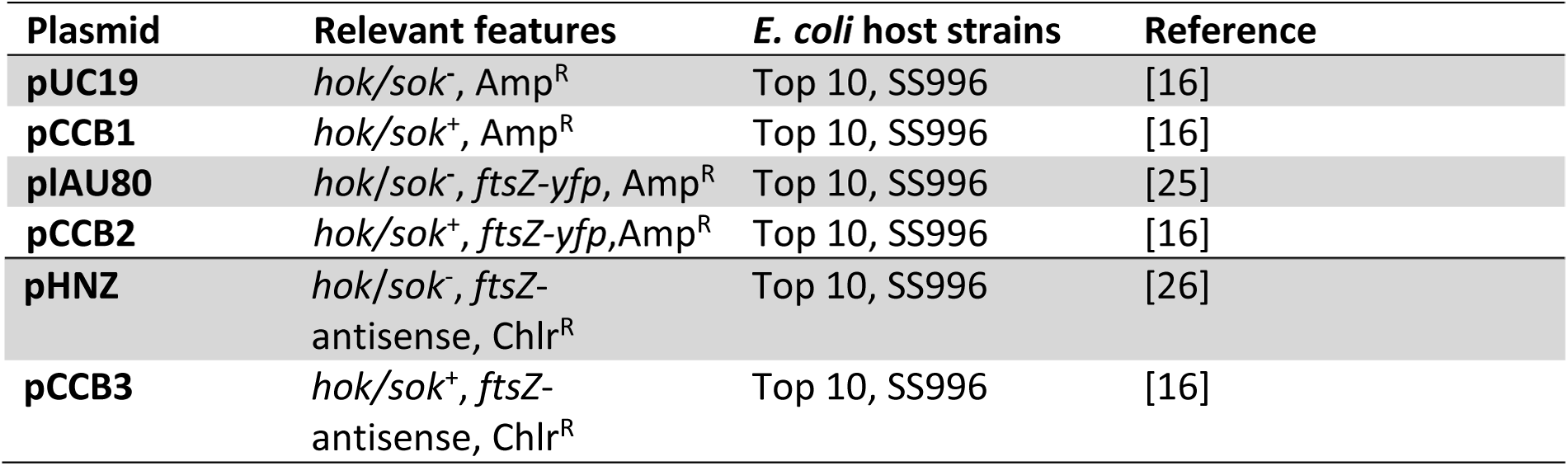
Plasmids and bacterial strains.

### 2.2 Preparation of competent cells

Host *E. coli* cells were made competent chemically (using Cacl_2_) for subsequent transformation with the plasmids indicated. A single colony of the host cell was grown in 5ml LB broth for 2-3hrs with vigorous shaking (≈250rpm), then transferred to 100ml of pre-warmed LB broth at 37°C. This was incubated at 37°C with vigorous shaking (250 rpm) to an optical density of 0.6 at 600nm (checked with spectrophotometer). Cells were chilled on ice for 15mins and transferred into 50ml sterile centrifuge tubes, which were then centrifuged at 2000 xg for 10mins, with temperature pre-set at 0°C. The pellets were then suspended in 15ml of sterile ice-cold 0.1M CaCl_2_ by gentle pipetting, cooled on ice for 15mins and centrifuged again for 10mins. They were again suspended in 4ml 0.1M CaCl_2_ in 15% glycerol and 200µl aliquots were stored at −80°C until used.

### 2.3 Plasmid extraction

Overnight cultures of the bacteria cells containing the desired plasmid were grown in LB media containing selective antibiotics appropriate for each plasmid in a Stuart orbital incubator S1500 at 37°C for 16hrs with shaking (180 rpm). Plasmids were extracted using QIAgene Miniprep kit (QIAGEN #27104), following the manufacturer’s protocol.

### 2.4 Transformation of bacteria with plasmids

The competent bacteria cells to be transformed and the plasmid were allowed to thaw gently on ice. 5µl of plasmid (≈5ng) was then added to 50µl of competent cells in 1.5ml microfuge tubes, mixed gently and incubated on ice for 30mins. The cells were shocked by placing the tube in a water bath at 42°C for 20-30sec. Cells were recovered by incubating with 500µl of SOC medium for 1-2hrs, and plating 50µl of the sample on LB agar plates containing appropriate antibiotics for selection. After overnight incubation, a single colony was streaked on agar plate containing the selective antibiotic for further growth. To further check the success of the transformation, plasmid extraction was carried out with the transformants using QIAprep miniprep kit and the resulting plasmid band checked against the size of the original plasmid by electrophoresis on 1% agarose gel. Glycerol stocks of transformants were stored at −80°C.

### 2.5 Growth and antibiotic susceptibility assays

Overnight bacterial cultures were diluted to about 1 ×10^6^ CFU (approximately 1 in 1000 dilution) in 96 well plates and grown as already described [16]. The antibiotic susceptibility assays were done by both disk diffusion antibiotic testing and broth dilution method. MIC was determined and scored as the lowest concentration of the drug at which no observable growth of the bacterial culture occurred after 18hrs of incubation.

### 2.6 Phase contrast microscopy

200µl samples were collected from bacterial cultures treated with sub-inhibitory concentration of doxycycline (0.5µM) and processed as earlier described [27]. The number of cells showing the ghost cell morphology was counted per field of view, and the average expressed as percentages in relation to the normal cell morphology.

## 3 Results

### 3.1 Effect of the hok/sok locus on the susceptibility of E. coli lab strain to doxycycline

Since the *hok*/*sok* toxin/antitoxin system (which works via RNA:RNA interactions) is often associated with antimicrobial resistance, and doxycycline has been shown to interact with and inhibit cleavage of dsRNAs, we tested the effect of doxycycline on the growth of *hok*/*sok* host bacteria cells (*hok*/*sok*^+^ cells) in comparison with the *hok*/*sok*^-^ cells. Using K12 Top 10 strains in which both plasmids are very stable without antibiotic selection, disk diffusion antibiotic susceptibility tests were done with doxycycline disks on Lb agar plates. The results showed that the zone of inhibition is increased in cells containing the *hok*/*sok* locus when compared to the *hok*/*sok*^-^ cells (Figure 2A), indicating that the *hok*/*sok*^+^ cells are more susceptible to doxycycline. To confirm this observed increase in susceptibility of the *hok*/*sok*^+^ cells and determine the extent to which the *hok*/*sok* locus affects doxycycline susceptibility, antibiotic susceptibility assay was also carried out in broth media using 96 well plates. The results also showed that the *hok*/*sok*^+^ cells were more susceptible to doxycycline than the *hok*/*sok*^-^ cells. Lower amount of doxycycline was needed to inhibit culture growth in cells containing the *hok*/*sok* locus than in those that do not contain the locus (Figure 2). Whereas 0.6µM doxycycline completely inhibited the growth of *hok*/*sok*^+^ cells, the *hok*/*sok*^-^ still showed measurable growth beyond 0.7µM doxycycline concentration. This suggests that even though the *hok*/*sok* confers antibiotic resistance to the host cells, they may on the other hand confer increased doxycycline susceptibility to the cells.

**Figure 2:**
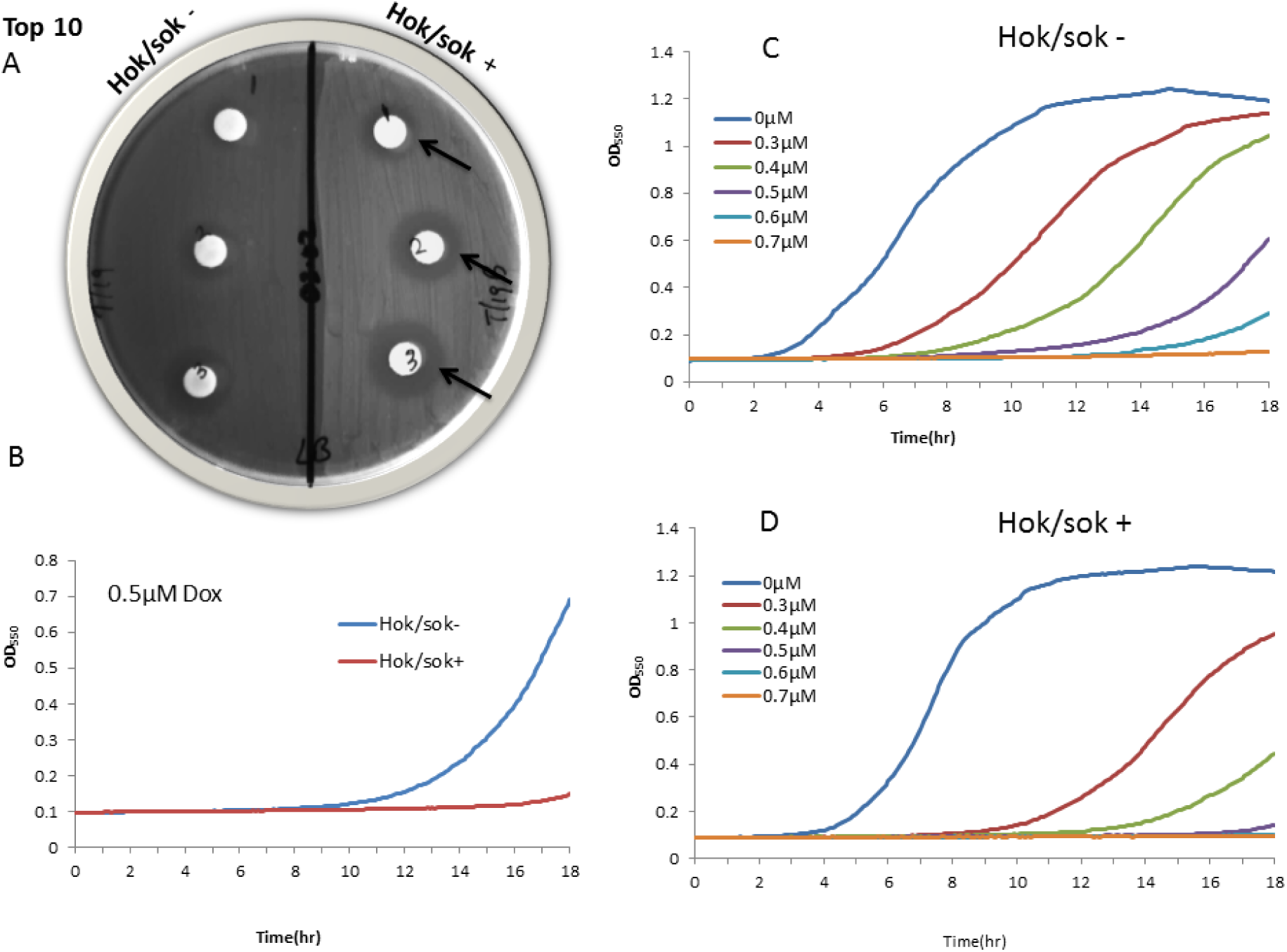
Effect of the *hok*/*sok* locus on *E. coli* lab strain (Top 10) susceptibility to doxycycline. Agar plate shows disk diffusion susceptibility tests of *E. coli* Top 10 strain, with increasing amounts of doxycycline applied to the disks (1, 2, 3=0.1, 0.2, 0.3µg doxycycline). Arrows indicate wider zones of inhibition in *hok*/sok^+^ cells compared to the control strain. Graphs show growth curves of cells in increasing concentrations of doxycycline, with the *hok*/*sok*^+^ cells showing higher growth inhibition. Data is representative of six repeat experiments.

### 3.2 Effect of the hok/sok locus on doxycycline susceptibility of SOS-negative E. coli strain

It has been established that the *hok*/*sok* locus acts as a stress response element in bacteria, and can functionally complement defective SOS response. Hence, we also explored the effect of the *hok*/*sok* locus on doxycycline susceptibility of an *E. coli* strain that is SOS-defective. This strain is ordinarily unable to withstand stressful growth conditions such as high temperature and antibiotic treatment, hence very limited growth in such conditions. However, with the *hok*/*sok* locus, it grows unrestricted in stressful growth conditions. Doxycycline susceptibility tests showed that the *hok*/*sok*^+^ cells were not only more susceptible, but the increase in susceptibility of the *hok*/*sok*^+^ cells to doxycycline is much more pronounced in this strain than in the lab strain (Figure 3). Whereas 1µM doxycycline did not inhibit the growth of *hok*/*sok*^-^ by half, the growth of the *hok*/*sok*^+^ was completely inhibited at the same drug concentration. This is quite the opposite of what was previously observed with some other antibiotics such as ampicillin and amoxicillin, in which case the *hok*/*sok*^-^ cells were more susceptible [16]. This suggests that the more pronounced the growth enhancing effects of the *hok*/*sok* locus in a strain is, the more it increases its susceptibility to doxycycline. In other words, the more operative/effective the *hok*/*sok* locus is at enhancing bacterial stress response and survivability, the more it increases the host cell susceptibility to doxycycline.

**Figure 3:**
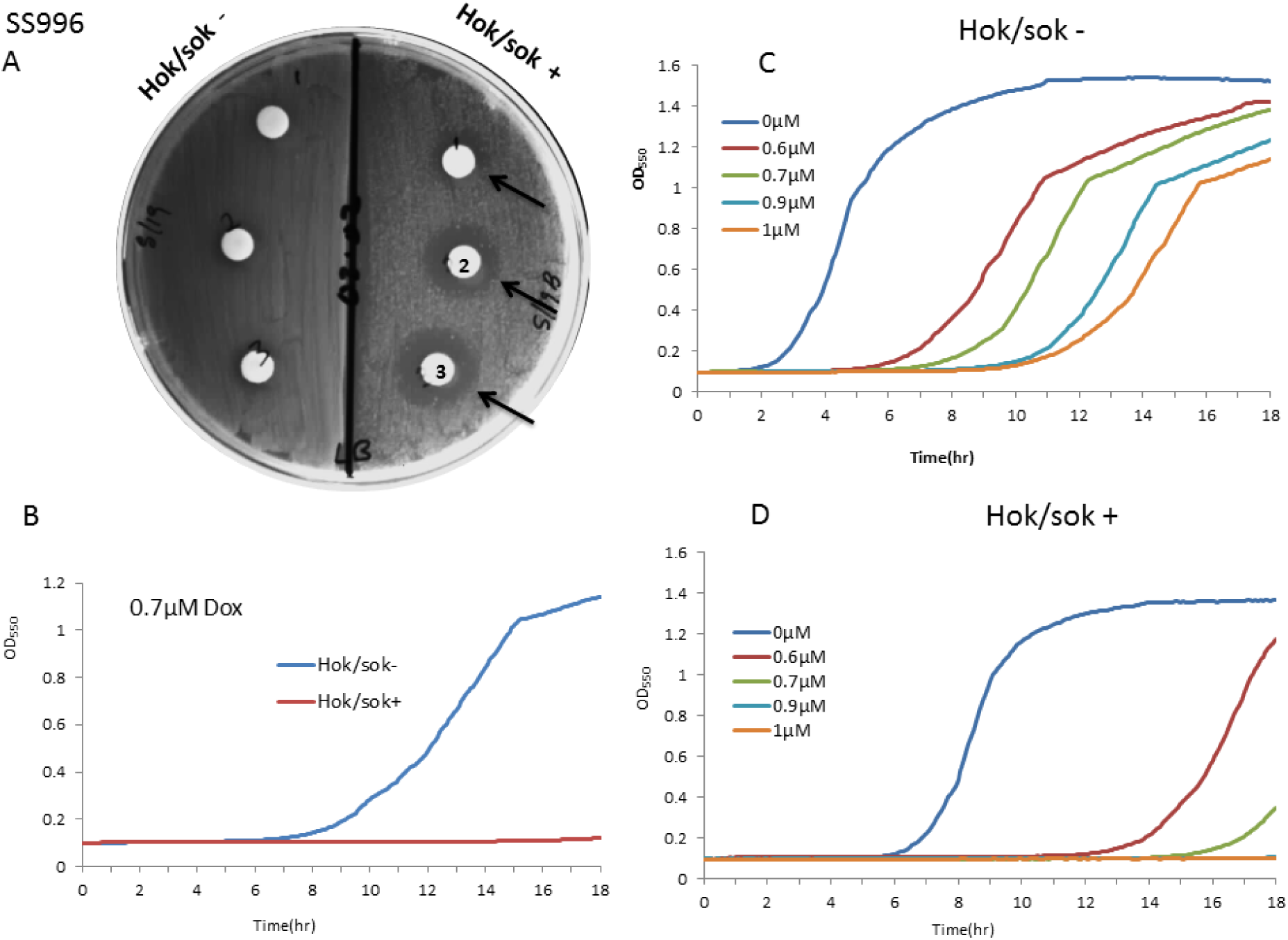
Effect of the *hok*/*sok* locus on the susceptibility of SOS-deficient *E. coli* (SS996) to doxycycline. Agar plate shows susceptibility tests of *E. coli* SOS=deficient strain (SS996) with increasing amounts of doxycycline applied to disks (1, 2, 3=0.1, 0.2, 0.3µg doxycycline). Arrows indicate wider zones of inhibition in *hok*/sok^+^ cells compared to the control strain. Graphs show growth curves of cells in increasing concentrations of doxycycline, with the *hok*/*sok*^+^ cells showing higher growth inhibition. Data is representative of six repeat experiments.

### 3.3 Effect of the hok/sok associated β-lactam resistance on doxycycline susceptibility

The *hok*/*sok* locus has also been shown to increase host cell tolerance/resistance to the β-lactam antibiotics (specifically ampicillin and amoxicillin), and this is more apparent in the SOS-deficient strain (SS996). Hence, we also explored the effect of this *hok*/*sok*-associated β-lactam resistance on the susceptibility of the cells to doxycycline. In Top 10 strain, the results showed that the *hok*/*sok*^+^ cells were still more susceptible to doxycycline than the *hok*/*sok*^-^ cells even in the presence of ampicillin (Figure 4). When the growth of the cells was compared in LB and LB amp media, there was no difference in the growth pattern and doxycycline susceptibility of the cells (D). This indicates that doxycycline susceptibility of the cells is not affected by the presence of ampicillin in the media (and the associated resistance), suggesting that the increase in doxycycline susceptibility in *hok*/*sok*^+^ cells occurs via a mechanism that is not antagonistic to the mechanism of ampicillin resistance.

**Figure 4:**
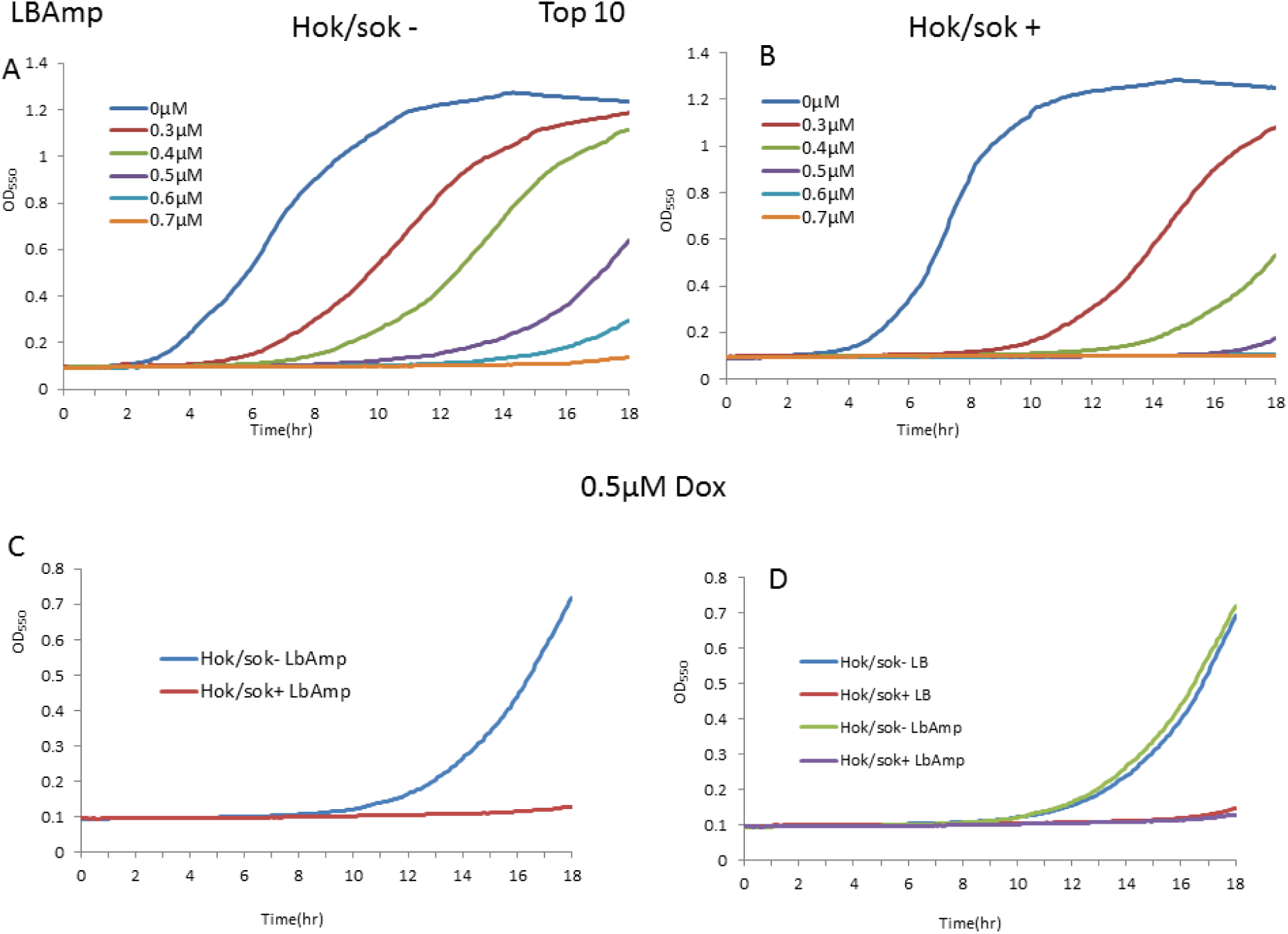
Effect of the hok/sok associated β-lactam resistance on doxycycline susceptibility. Graphs show growth curves of Top 10 cells in increasing concentrations of doxycycline in LbAmp media (containing 100 µg/ml ampicillin), with the *hok*/*sok*^+^ cells showing higher growth inhibition (C). Doxycycline susceptibility of the *hok*/*sok*^+^ cells remains the same in both Lb and LbAmp media (D). Data is representative of six repeat experiments.

In the SS996 strain, there was also no difference in the growth pattern and doxycycline susceptibility of the *hok*/*sok*^+^ cells when compared in LB and LB amp media (Figure 5). This further indicates that doxycycline susceptibility of the *hok*/*sok*^+^ cells is not affected by the associated β-lactam (ampicillin) resistance. However, the control (*hok*/*sok*^-^) cells showed greater growth inhibition and improved susceptibility to doxycycline in LbAmp media (because it is more susceptible to ampicillin as has been earlier reported). These results indicate that doxycycline can overcome the ampicillin resistance imparted by the *hok*/*sok* locus to host cells, and that both doxycycline and ampicillin can work synergistically to enhance bacterial killing/growth inhibition.

**Figure 5:**
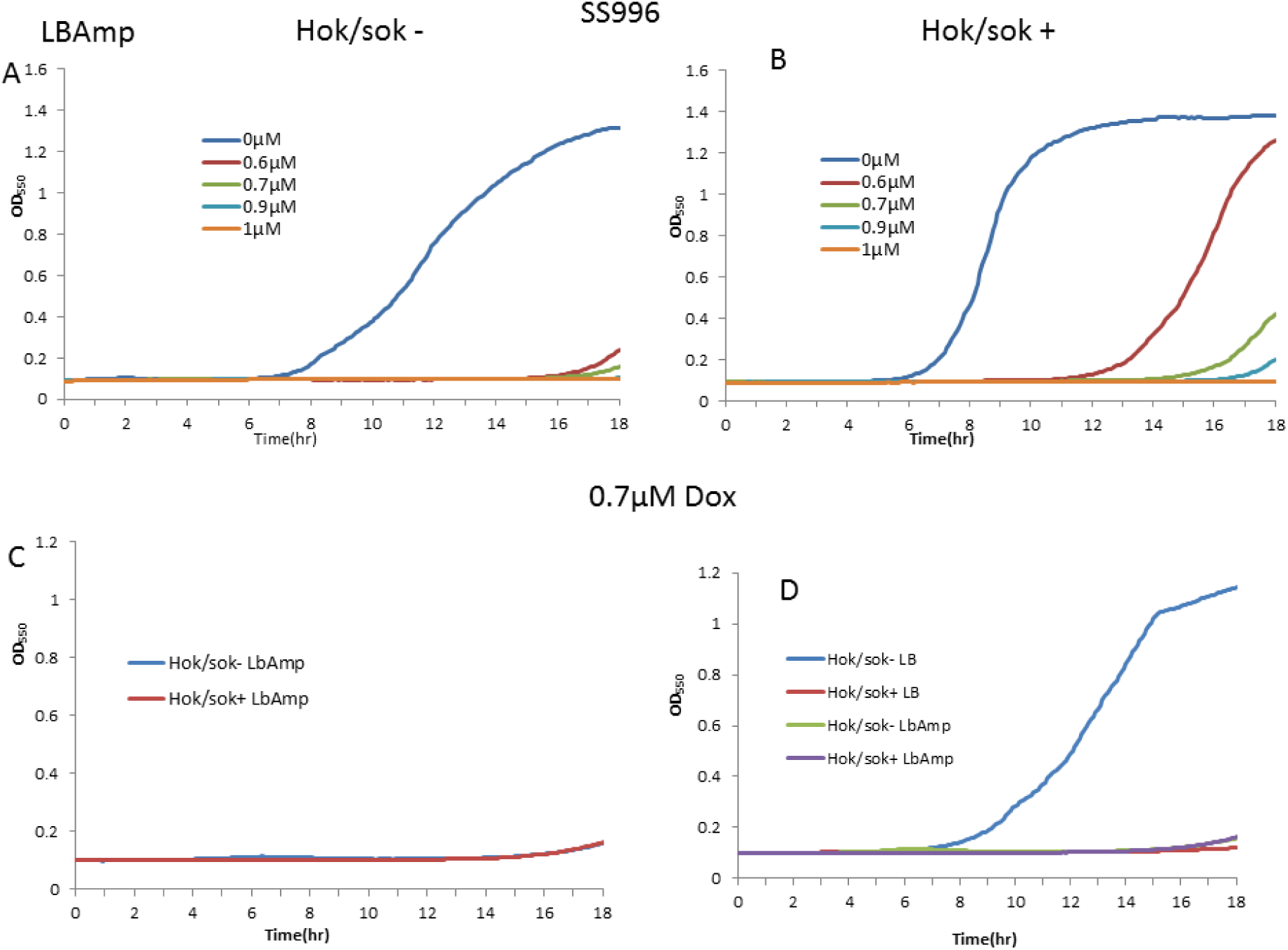
Effect of the hok/sok associated β-lactam resistance on doxycycline susceptibility. Graphs show growth curves of SS996 cells in increasing concentrations of doxycycline in LbAmp media (containing 100 µg/ml ampicillin). Doxycycline susceptibility of the *hok*/*sok*^+^ cells remains the same in both Lb and LbAmp media (D), but the *hok*/*sok*^-^ also show increased susceptibility in LbAmp media. Data is representative of six repeat experiments.

### 3.4 MIC of doxycycline in hok/sok host cells

MIC determination for doxycycline also showed that lower amounts of doxycycline are required to completely inhibit the growth of cells containing the *hok*/*sok* than those that do not contain the *hok*/*sok* locus (Table 2).

**Table 2:**
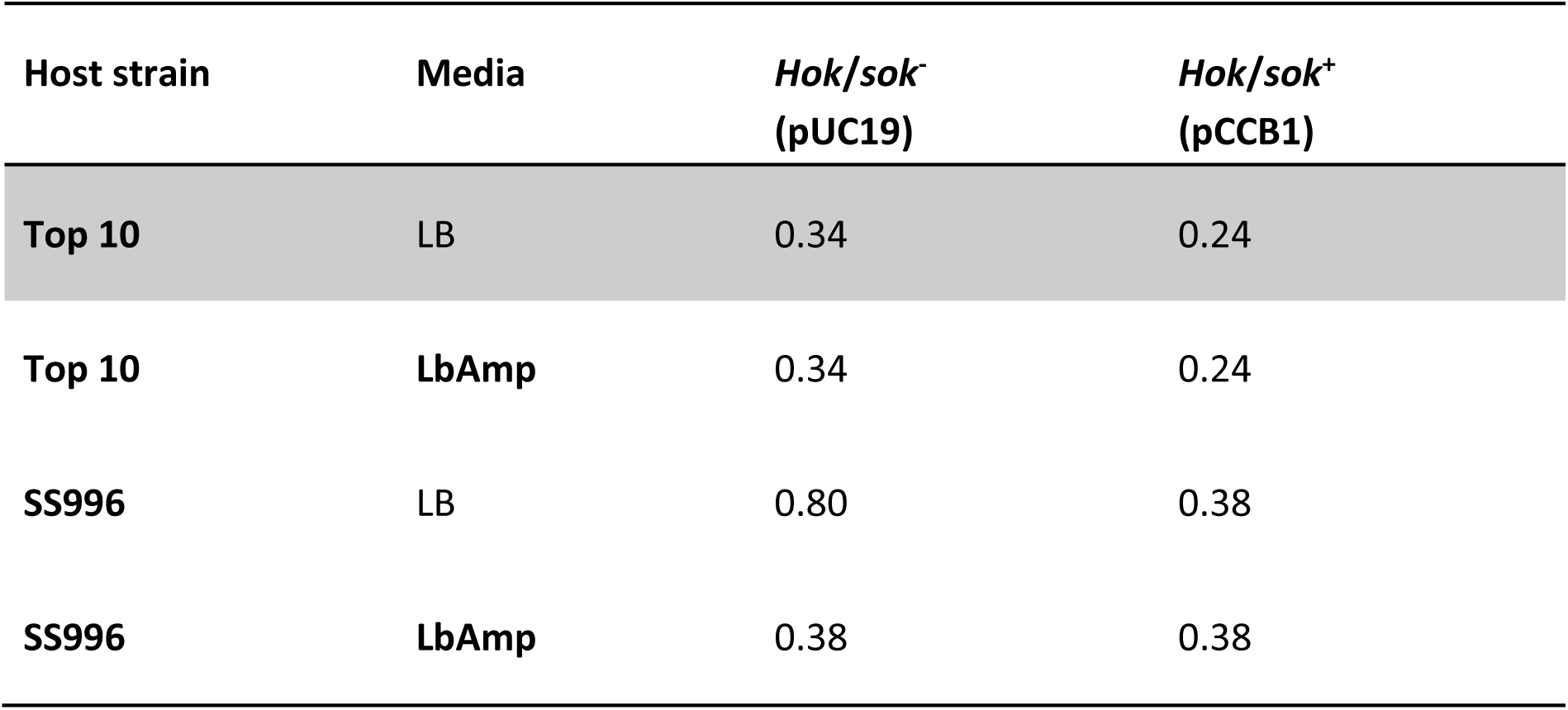
MIC (µg/ml) of doxycycline in *E. coli* cells containing plasmid-borne hok/sok locus.

In order to substantiate that the observed increase in doxycycline susceptibility is brought about by the *hok*/*sok* locus, we also tested the susceptibility of cells containing two other *hok*/*sok* plasmids (pCCB2 and pCCB3) and their control cells (plAU80 and pHNZ) to doxycycline. The results still showed increased susceptibility of the *hok*/*sok*^+^ cells to doxycycline (Figures 6 and 7). This further asserts that the *hok*/*sok* locus enhances host cell susceptibility to doxycycline.

**Figure 6:**
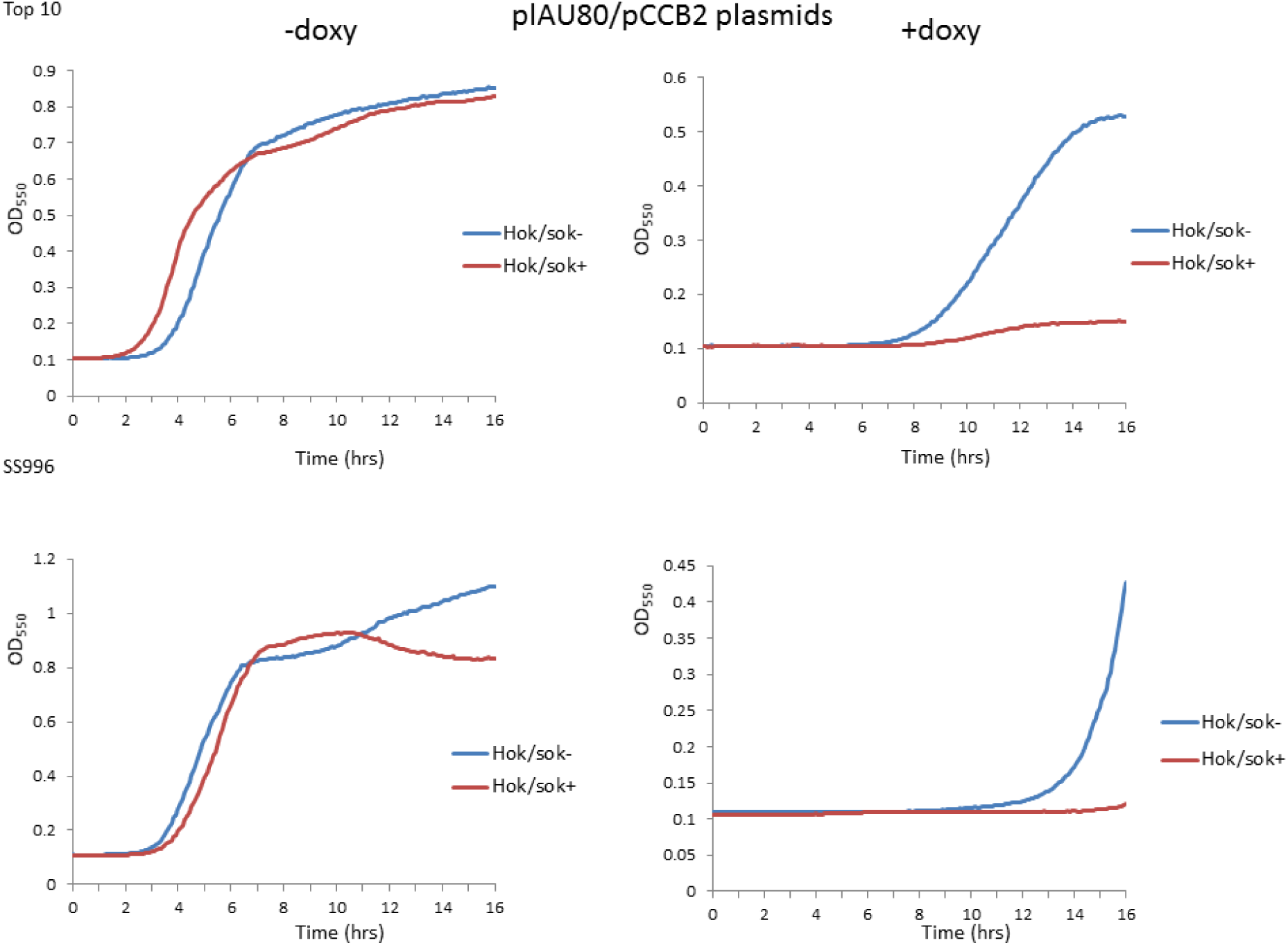
Effect of the *hok*/*sok* locus on doxycycline susceptibility. Graphs show growth curves of Top 10 and SS996 cells containing pCCB2 *hok*/*sok* plasmids and controls treated with 1µM doxycycline, with the *hok*/*sok*^+^ cells showing higher growth inhibition.

**Figure 7:**
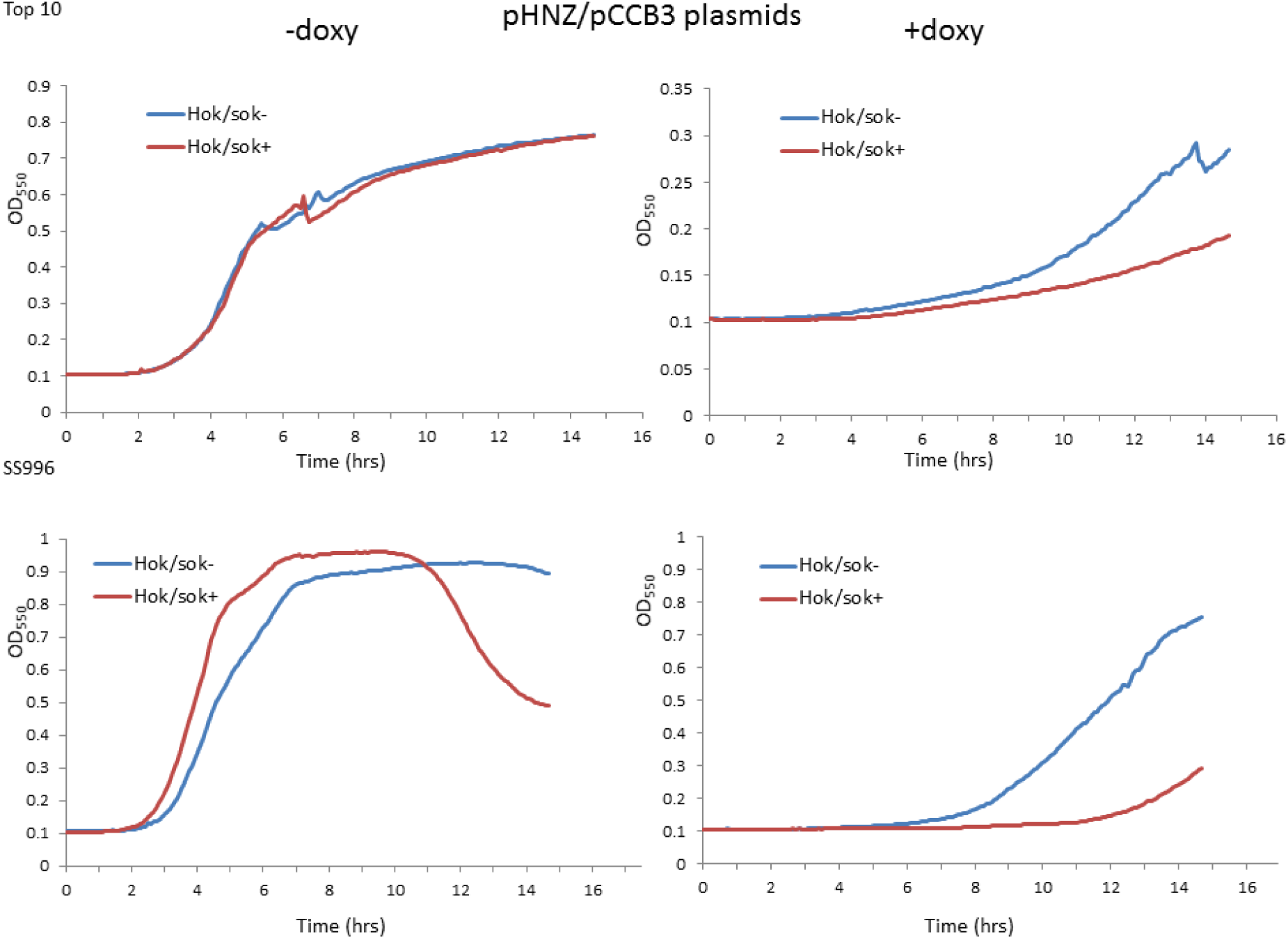
Effect of the *hok*/*sok* locus on doxycycline susceptibility. Graphs show growth curves of Top 10 and SS996 cells containing pCCB3 *hok*/*sok* plasmids and controls treated with 1µM doxycycline, with the *hok*/*sok*^+^ cells showing higher growth inhibition.

### 3.5 Effect of doxycycline on the morphology of hok/sok host cells

The *hok*/*sok* toxin/antitoxin system induces host cell death via a post-segregational killing mechanism that is associated with a characteristic cell morphology known as “ghost cells” due to the activity of the Hok toxin. In order to ascertain whether the observed increase in doxycycline susceptibility of *hok*/*sok* host cells is connected to the toxin/antitoxin system, we examined the morphology of cells treated with sub-inhibitory concentration of doxycycline (0.5µM ≈0.2µg/ml) by phase contrast microscopy to see whether the cells show the typical morphology of Hok toxin activity/cell death in bacteria (ghost cells). The images obtained show increased number of the *hok*/*sok*^+^ cells with morphologic changes suggestive of Hok toxin activity (ghost cells). These cells appear denser at the poles than in the middle, indicating impaired membrane permeability associated with Hok killing (Figure 8). About 70% of the *hok*/*sok*^+^ cells showed ghost cell morphology, whereas less than 10% of the *hok*/*sok*^-^ cells showed a similar morphology (similar to the untreated cells). This suggests that the Hok toxin is activated in bacteria cells containing the *hok*/*sok* locus following doxycycline treatment.

**Figure 8:**
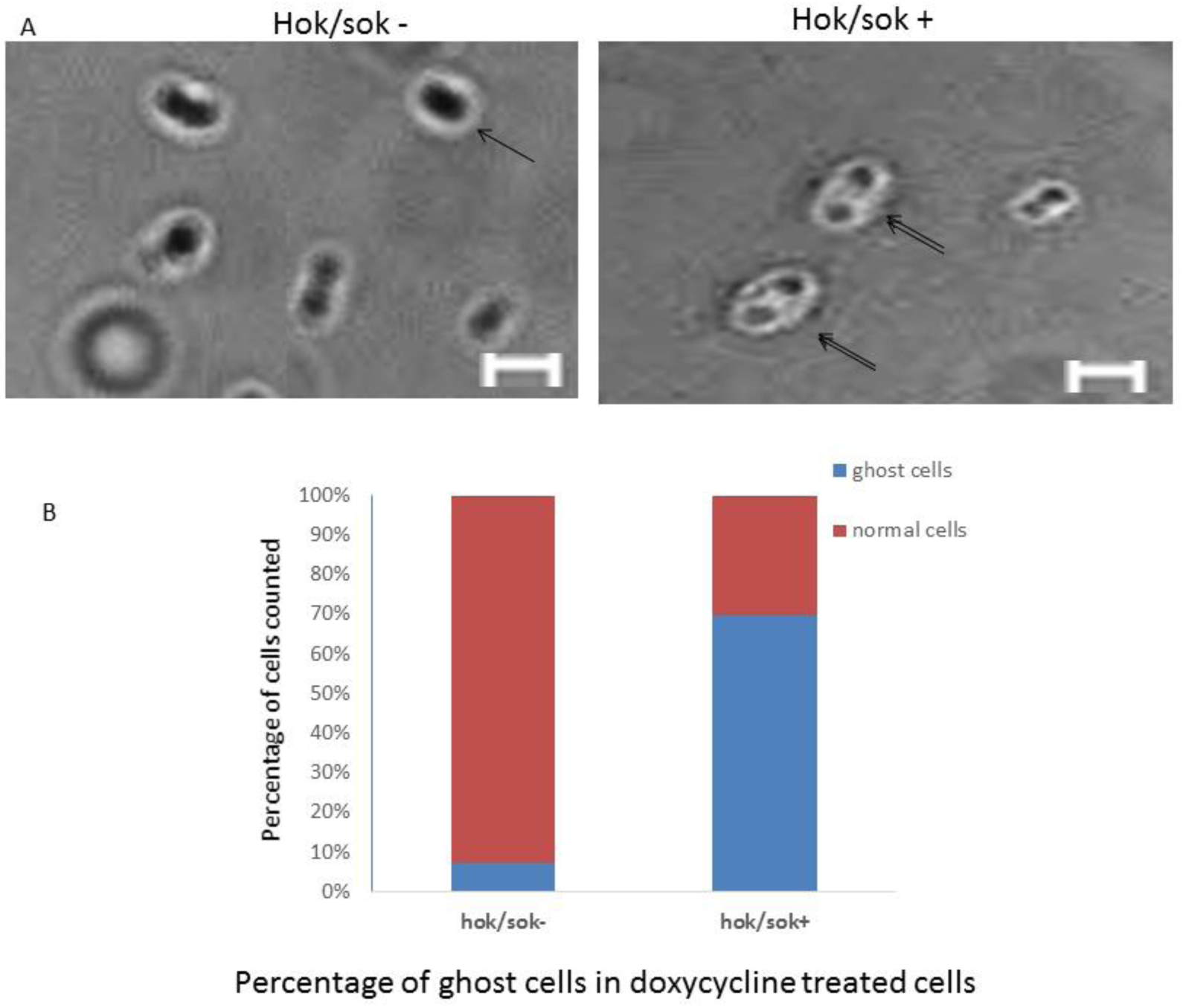
Phase contrast microscopy images *hok*/*sok*^+^ and *hok*/*sok*^−^ cells treated with sub-inhibitory concentration of doxycycline (0.5µM). Double arrows indicate cells that appear denser at the poles than the middle (“ghost” cells), which are morphological changes associated with Hok toxin activity, compared to normal cells (indicated by single arrow). Scale bar= 2µm. Histogram shows the percentage of ghost cells to normal cells counted from an equal area of the phase contrast microscopy images of *hok*/*sok*^+^ and *hok*/*sok*^-^ bacteria samples, with about 70% ghost cells in the *hok*/*sok*^+^ sample and about 10% ghost cells in the *hok*/*sok*^-^ sample.

## 4 Discussion

The *hok*/*sok* locus has been shown to contribute to bacterial stress response by helping the cells survive adverse growth condition such as high temperature and antibiotic treatment [16]. Particularly, it confers a selective advantage to host bacteria cells in ampicillin-selective media, thereby enhancing ampicillin (and β-lactam antibiotics) resistance. This effect is especially apparent in strains that are defective in the SOS response (such as SS996), and has been associated with alterations in FtsZ activity [27]. It has also been shown to be closely associated with plasmids encoding ESBLs, especially CTX-M beta-lactamases [20], and many other antibiotics resistance genes [17-19].

The *hok*/*sok* locus contributes to β-lactam resistance in two ways: by increasing the tolerance of host cells to higher amounts of the drug [16], and by ensuring the stabilization/maintenance and propagation of the resistance genes normally carried on the *hok*/*sok* plasmids [15, 28]. The plasmid stabilization/maintenance function of the *hok*/*sok* locus is a very useful tool that ensures host bacteria evade multiple antibiotic treatment, thus sustaining and spreading antibiotic resistance. This combined/multiple mechanisms of antibiotic resistance makes the *hok*/*sok* gene critical in the fight against antibiotic resistance. With such contributions in enhancing antibiotic tolerance and resistance gene propagation, the *hok*/*sok* locus cannot be overlooked in mapping out strategies to combat antibiotic resistance. Hence, the observations of increased *hok*/*sok* host cell susceptibility to doxycycline in this study could open up opportunities to fight antibiotic resistance using the tools supplied by the bacteria themselves. In other words, the *hok*/*sok* locus presents a potential drug target to combat antibiotic resistance and propagation of resistance elements.

Considering the fact that the antibiotic usefulness of the tetracyclines has been greatly limited due to the development of resistance, it is particularly interesting to observe that it could be useful for killing cells that are multidrug resistant when associated with the *hok*/*sok* locus. When the doxycycline susceptibility of the *hok*/*sok* host cells was compared in media with and without ampicillin, there was no difference in the growth pattern of the cells. This indicates that the *hok*/*sok* locus renders bacteria more susceptible to doxycycline irrespective of resistance to other antibiotics, suggesting that the *hok*/*sok*–associated increase in doxycycline susceptibility occurs via a mechanism that is not antagonistic to the mechanism of ampicillin resistance. Hence, doxycycline may be a more effective therapy against infections by pathogens containing the hok/sok locus, particularly in cases of suspected beta-lactam resistance. Again, the synergistic action of doxycycline with ampicillin may present a better therapeutic combination in cases where the genetic makeup of the cells cannot be determined.

The observed morphologic appearance of the *hok*/*sok* host cells treated with doxycycline suggests that *hok*/*sok* locus enhances doxycycline susceptibility by inducing Hok toxin expression and killing of the cells. Since doxycycline inhibits RNAse III cleavage and processing of dsRNAs [23, 24], it would inhibit RNAse III degradation of the *hok* mRNA:Sok RNA complex leading to the eventual decay of the labile Sok RNA and releasing the *hok* mRNA for translation and subsequent cell killing. Activation of Hok toxin activity/self-killing in bacteria containing *hok*/*sok* plasmids has also been reported by Faridani et al (2006) using peptide nucleic acids (PNA) that target Sok RNA [22]. This suggests that antimicrobial agents that could inhibit RNAse degradation of the *hok*/*sok* dsRNA complex (such as RNA ligands) could therefore be exploited as a therapeutic strategy to induce self-killing in the cells and enhance antibiotic efficacy, as indicated here with doxycycline.

## References

1. Erdmann, V.A., et al., Regulatory RNAs. Cellular and Molecular Life Sciences, 2001. 58(7): p. 960–977.

2. Waters, L.S. and G. Storz, Regulatory RNAs in Bacteria. Cell, 2009. 136(4): p. 615–628.

3. Wagner, E.G.H. and R.W. Simons, Antisense RNA Control in Bacteria, Phages, and Plasmids. Annual Review of Microbiology, 1994. 48(1): p. 713–742.

4. Gerdes, K. and E.G.H. Wagner, RNA antitoxins. Current Opinion in Microbiology, 2007. 10(2): p. 117–124.

5. Fozo, E.M., M.R. Hemm, and G. Storz, Small Toxic Proteins and the Antisense RNAs That Repress Them. Microbiology and Molecular Biology Reviews, 2008. 72(4): p. 579–589.

6. Fozo, E.M., et al., Abundance of type I toxinâ€”antitoxin systems in bacteria: searches for new candidates and discovery of novel families. Nucleic Acids Research. 38(11): p. 3743–3759.

7. Thisted, T. and K. Gerdes, Mechanism of post-segregational killing by the hok/sok system of plasmid R1. Sok antisense RNA regulates hok gene expression indirectly through the overlapping mok gene. J Mol Biol, 1992. 223(1): p. 41–54.

8. Gerdes, K., et al., Mechanism of killer gene activation. Antisense RNA-dependent RNase III cleavage ensures rapid turn-over of the stable Hok, SrnB and PndA effector messenger RNAs. Journal of Molecular Biology, 1992. 226(3): p. 637–649.

9. K, G., et al., Translational control and differential RNA decay are key elements regulating postsegregational expression of the killer protein encoded by the parB locus of plasmid R1. J. Mol. Biol., 1988. 203: p. 119.

10. Gerdes, K., T. Thisted, and J. Martinussen, Mechanism of post-segregational killing by the hok/sok system of plasmid R1: sok antisense RNA regulates formation of a hok mRNA species correlated with killing of plasmid-free cells. Mol Microbiol, 1990. 4(11): p. 1807–18.

11. Gerdes, K., et al., Antisense RNA-regulated programmed cell death. Annu Rev Genet, 1997. 31: p. 1–31.

12. Franch, T., A.P. Gultyaev, and K. Gerdes, Programmed cell death by hok/sok of plasmid R1: processing at the hok mRNA 3’-end triggers structural rearrangements that allow translation and antisense RNA binding. J Mol Biol, 1997. 273(1): p. 38–51.

13. K, G., R. PB, and M. S, Unique type of plasmid maintenance function: postsegregational killing of plasmid free cells. Proc. Natl. Acad. Sci. USA, 1986. 83: p. 3116.

14. K, G., L. Jel, and M. S, Stable inheritance of plasmid R1 requires two different loci. J. Bacteriol., 1985. 161: p. 292.

15. Gerdes, K., J.S. Jacobsen, and T. Franch, Plasmid stabilization by post-segregational killing. Genet Eng (N Y), 1997. 19: p. 49–61.

16. Chukwudi, C.U. and L. Good, The role of the hok/sok locus in bacterial response to stressful growth conditions. Microb Pathog, 2015. 79: p. 70–9.

17. Campbell, C.S. and R.D. Mullins, In vivo visualization of type II plasmid segregation: bacterial actin filaments pushing plasmids. The Journal of Cell Biology, 2007. 179(5): p. 1059–1066.

18. Blohm, D. and W. Goebel, Restriction map of the antibiotic resistance plasmid R1drd-19 and its derivatives pKN102 (R1drd-19B2) and R1drd-16 for the enzymes BamHI, HindIII, EcoRI and SalI. Molecular and General Genetics MGG, 1978. 167(2): p. 119–127.

19. Clerget, M., M. Chandler, and L. Caro, The structure of R1drd19: A revised physical map of the plasmid. Molecular and General Genetics MGG, 1981. 181(2): p. 183–191.

20. Mnif, B., et al., Molecular characterization of addiction systems of plasmids encoding extended-spectrum beta-lactamases in Escherichia coli. J Antimicrob Chemother, 2010. 65(8): p. 1599–603.

21. Pedersen, K. and K. Gerdes, Multiple hok genes on the chromosome of Escherichia coli. Molecular Microbiology, 1999. 32(5): p. 1090–1102.

22. Faridani, O.R., et al., Competitive inhibition of natural antisense Sok-RNA interactions activates Hok-mediated cell killing in Escherichia coli. Nucleic Acids Res, 2006. 34(20): p. 5915–22.

23. Chukwudi, C.U. and L. Good, Interaction of the tetracyclines with double-stranded RNAs of random base sequence: new perspectives on the target and mechanism of action. J Antibiot (Tokyo), 2016. 69(8): p. 622–30.

24. Chukwudi, C.U. and L. Good, Doxycycline inhibits pre-rRNA processing and mature rRNA formation in E. coli. J Antibiot (Tokyo), 2019.

25. Ebersbach, G., et al., Novel coiled-coil cell division factor ZapB stimulates Z ring assembly and cell division. Mol Microbiol, 2008. 68(3): p. 720–35.

26. Goh, S., et al., Concurrent growth rate and transcript analyses reveal essential gene stringency in Escherichia coli. PLoS ONE, 2009. 4(6): p. e6061.

27. Chukwudi, C.U. and L. Good, Phenotypic indications of FtsZ inhibition in hok/sok-induced bacterial growth changes and stress response. Microb Pathog, 2018. 114: p. 393–401.

28. Gerdes, K., The parB (hok/sok) Locus of Plasmid R1: A General Purpose Plasmid Stabilization System. Nat Biotech, 1988. 6(12): p. 1402–1405.

